# RNA-seq transcriptome analysis reveals LTR-retrotransposons modulation in Human Peripheral Blood Mononuclear Cells (PBMCs) after *in vivo* Lipopolysaccharides (LPS) injection

**DOI:** 10.1101/2019.12.20.884494

**Authors:** Maria Paola Pisano, Olivier Tabone, Maxime Bodinier, Nicole Grandi, Julien Textoris, François Mallet, Enzo Tramontano

## Abstract

Human Endogenous Retroviruses (HERVs) and Mammalian apparent LTR-retrotransposons (MaLRs) are retroviral sequences that integrated into the germline cells millions year ago. Transcripts of these LTR-retrotransposons are present in several tissues, and their expression is modulated in pathological conditions, although their function remains often far from being understood. In this work, we focused on the HERVs/MaLRs expression and modulation in a scenario of immune system activation. We used a public dataset of Human Peripheral Blood Mononuclear Cells (PBMCs) RNA-seq from 15 healthy participants to a clinical trial before and after the exposure to Lipopolysaccharide (LPS), for which we established an RNA-seq workflow for the identification of expressed and modulated cellular genes and LTR-retrotransposon elements.

**Importance:** We described the HERV and MaLR transcriptome in PBMCs, finding that about 8.4 % of the LTR-retrotransposons loci were expressed, and identifying the betaretrovirus-like HERVs as those with the highest percentage of expressed loci. We found 4,607 HERVs and MaLRs loci that were modulated as a result of *in vivo* stimulation with LPS. The HERV-H group showed the highest number of differentially expressed most intact proviruses. We characterized the HERV and MaLR loci differentially expressed checking their genomic context of insertion and, interestingly, we found a general co-localization with genes that are involved and modulated in the immune response, as consequence of LPS stimulation. The analyses of HERVs and MaLRs expression and modulation show that this LTR-retrotransposons are expressed in PBMCs and regulated in inflammatory settings. The similar regulation of HERVs/MaLRs and genes after LPS stimulation suggests possible interactions of LTR-retrotransposons and the immune host response.

## Background

A large proportion of the human genome consists of repeated elements, including Human Endogenous Retroviruses (HERVs) and Mammalian Apparent LTRs Retrotransposons (MaLRs) (1). These elements are remnants of ancestral and independent infections within the germline cells that took place million years ago (2). At the time of integration, the HERV genome was initially composed of four retroviral genes (*gag, pro, pol*, and *env*) flanked by two Long Terminal Repeats (LTRs). Similarly, the genomic structure of MaLRs, more ancient retroviral elements, is similar but characterized by the absence of the *env* gene (3, 4). During the time, these elements have mostly accumulated abundant mutations often compromising their coding capability, and a great number of solitary LTRs were obtained by recombination occurrences (2, 4), as clearly observably when comparing human LTR-retrotransposons integrations with their orthologs in primates (5, 6). The potential involvement of these elements in the human biology has been investigated in several studies (7) which have shown the presence of few preserved retroviral Open Reading Frames (ORFs) expressed in human tissues (8–10), including Syncytin-1, a functional HERV-W envelope protein that is involved into the trophoblastic-cells fusion in placenta development (11, 12). Another important function of HERVs and MaLRs insertions is linked to species evolutionary developments. For example, data suggest that the high density of HERV integrations had a role in promoting and maintaining the diversity of multigenic regions, as in the case of human Major Histocompatibility Complex (MCH) (13–15). Furthermore, the promoters, enhancers and poyadenylation signals within the LTR sequence allow the proviral and solitary LTRs to play a role into the expression of the host genes (16). More specifically, the integration of newly provided regulatory signals at one of the ends of the host gene sequence could result in the modulation of its expression (16), while the presence of viral promoters or polyadenylation signals near to the gene or within its intron could result in altered RNA splicing (16–19). In this respect, the HERVs and MaLRs contribution in shaping and influencing the human innate immunity is an argument of particular interest (20). Indeed, in some cases LTR-retrotransposon derived antigens could be recognized as pathogen-associated molecular patterns (PAMPs) by Pattern Recognition Receptors (PRRs) such as the transmembrane proteins Toll-Like Receptors (TLRs) (21, 22). In these cases, the activation of PRRs evokes complex cellular signaling pathways altering gene expression for transducing pro-inflammatory signals (20). Despite the fact that on the one hand these interactions with PRRs are hypothesized to be positively involved in shaping the evolution of the immune response (21, 23), on the other hand, these mechanisms have been investigated for their possible contribution to the development of autoimmunity and inflammatory diseases (9, 24–26), like multiple sclerosis (27, 28). Interestingly, the activation of immune response through treatments with LPS or TNF-α can lead to an increase of HERVs and MaLRs expression (29, 30). For instance, a recent microarray-based study revealed the *in vitro* modulation of HERV/MaLR expression in PBMCs, after high- and low-dose LPS and Interferon-γ (IFN-γ) stimulations (29). A similar approach allowed to observe HERV/MaLR *in vivo* modulation in samples of blood in various contexts of injuries, also introducing a possible role of HERVs/MaLRs close to immunity-related genes in the regulation of their expression (31). In any case, many intriguing questions about the effective role of HERV and MaLR expression in immunity are still unsolved, and, in this respect, identifying the individual LTR-retrotransposon genomic localization and coding capacity could help to understand of their potential effects in this context (32). In this context, with the aim of investigating possible interactions between LTR-retrotransposons and the immune host response, we performed a comprehensive description of HERV and MaLR transcriptome observed in the analysis of a public RNA-seq dataset of PBMCs samples from patients *in vivo* stimulated with LPS. This experimental design has allowed mimicking an inflammation gram-negative induced. We focused our attention on the identification of specific expressed loci, as well as of those that were modulated as a result of stimulation with LPS. At last, we checked for similar regulation of HERVs/MaLRs and genes expression.

## Results

### Description of the LTR-retrotransposon transcriptome in PBMCs

With the aim to detect the HERV/MaLR transcriptome in PBMCs, we defined an RNA-seq pipeline to be used on a public RNA-seq (GEO:GSE87290) that included the PBMC transcriptome of 15 healthy volunteers before and after *in vivo* stimulation with 1ng/Kg LPS. Using this pipeline we could discriminate 424,515 loci from the *hervgdb4* database, including those of 197,341 HERVs and those of 227,174 MaLRs (33). Importantly, as the *hervgdb4* database has been created as part of the design of Affymetrix HERV-V3 array probes (33), the HERV and MaLR loci are included in the database as 881,603 *hervgdb4* fragments (Fig. S1) (33). Moreover, according to the annotation accuracy of the fragments and to their source, the loci were part of two different subsets, i) the highly informative HERV_prototypes and ii) the roughly annotated HERV_Dfam and MaLR_Dfam both collected from the Dfam database (Fig. S1) (34). Our analysis showed that 18,633 HERV *hervgdb4* loci and 17,053 MaLR *hervgdb4* loci were expressed in either stimulated and un-stimulated PBMCs samples, for a total of 35,686 loci, accounting for ∼ 8.4% of the LTR-retrotransposons in the human genome (Fig. S1). The majority of the LTR-retrotransposons expressed were part of the roughly-annotated HERV_Dfam and MaLR_Dfam subset, for which no information about the groups was available. Among the well-annotated HERV_prototypes we found expressed, 2084 were members of the class I gamma-like, 527 of the class II beta-like and 310 of the class III spuma/epsilon-like (Fig. 1a). When considering only the absolute number of expressed loci, just ahead form the HERV-H group, the PRIMA41 group was the one most represented. Instead, when considering the proportion of expressed loci among the HERV_prototypes groups, the most recently integrated group, HML-2, was the one showing the highest transcriptional activity, with more than 30% of its loci expressed. HML-10, HML-8 and HML-9 groups were also very active, with a proportion of 25.8%, 26.4% and 25% expressed loci, respectively. In general, the II beta-like groups were those showing overall the greatest percentage of activation. The proportions of expressed proviruses compared to solitary LTRs within the 3 classes were 55%, 61% and 64%, respectively (data not shown). Furthermore, in order to better define the HERV transcriptome in PBMCs, we decided to analyze the most intact proviruses that have been identified, classified and characterized in Vargiu *et al*. (4). These sequences have been collected by Vargiu *et al*. after the analysis of the software RetroTector (ReTe) on the human genome assembly GRCh 37/hg19, which searched for the presence of conserved motif hits and from these allowed to reconstruct the coordinates of 3173 HERVs satisfying distance constraints (4, 35). We identified 723 expressed ReTe proviruses and also in this case we found a large proportion of expressed beta-like elements. HML-4 (6 out of 12 ReTe proviruses) and HML-2 (43 out of 92 ReTe proviruses) were the most active groups (Fig. 1b). The group with the highest number of expressed ReTe proviruses was the class I gamma-like HERV-H, with 241 loci, representing the 23% of the whole group.

**Fig. 1.**
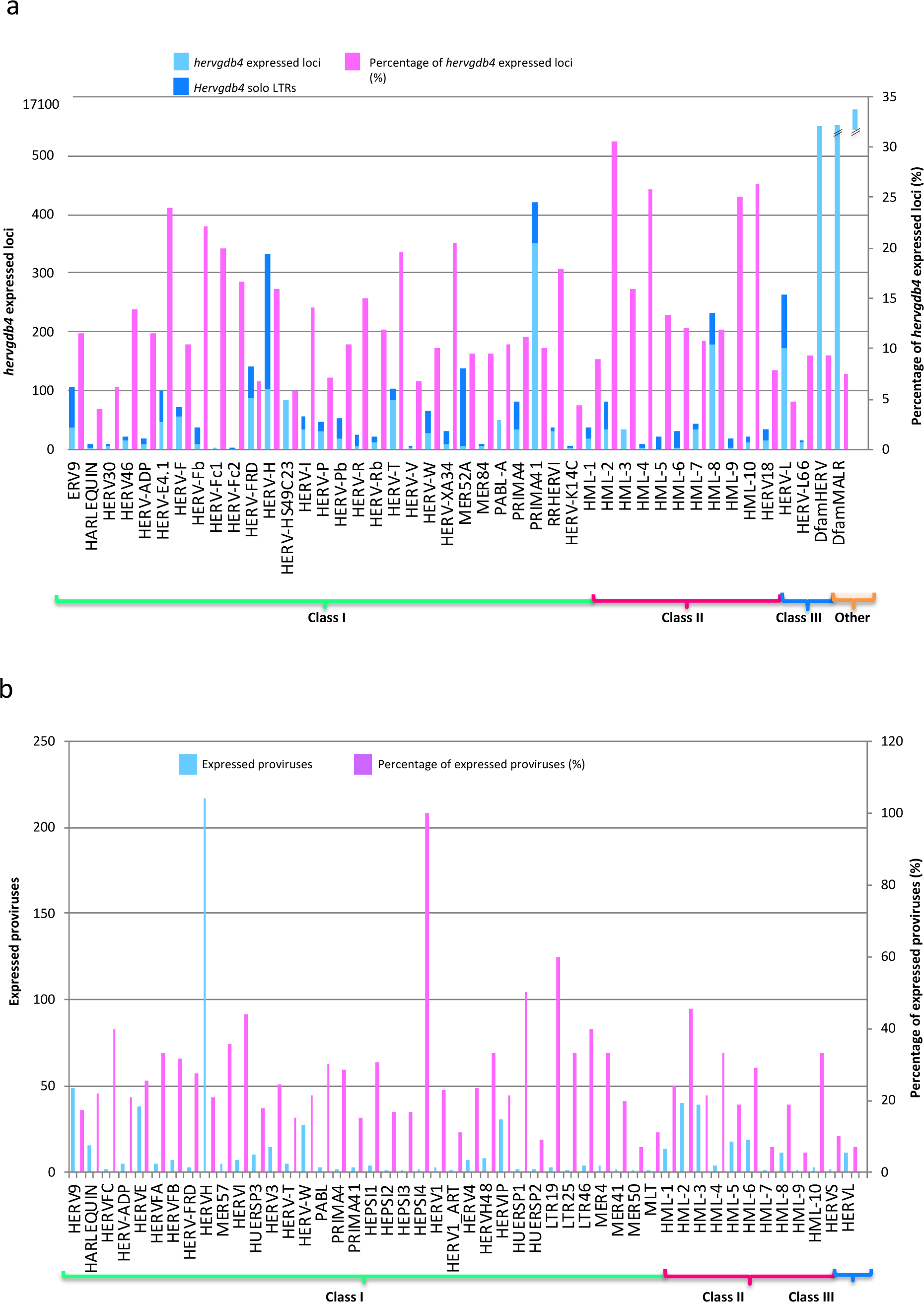
The LTR-retrotransposon transcriptome in PBMCs. Basal expression of the *hervgdb4* loci (a) and of the mostly intact HERV loci reported in Vargiu *et al*. (4) (b) in PBMCs. All the expressed elements are grouped by retroviral classes. and groups.

### Distinct transcriptional patterns induced by LPS stimulation

Next we wanted to assess the variability across the samples and to detect specific signatures induced by LPS stimulation. Hence, we analyzed the expression data of *hervgdb4* fragments in the 30 PBMC samples using the unsupervised Principal Component Analysis (PCA) (Fig. 2). The first Principal Component (PC1) explained the 49% of the variance across samples and revealed clustering specifically related to LPS response, dividing almost the totality of LPS-stimulated from not LPS-stimulated samples. Indeed, these data suggested that LPS response is the principal determinant for defining the inter-sample variability, showing *hervgdb4* fragments activation pattern specific of each of the two conditions. Moreover, we further investigated the transcriptional signature of the samples by performing hierarchical clustering on the 1,000 *hervgdb4* fragments with the highest mean of reads counts across samples (Fig. 3a). Results showed clear differences in expression of the *hervgdb4* fragments across samples, related to LPS stimulation. However, both PCA and hierarchical clustering analyses showed that the LTR-retrotransposon expression profile of samples 7, 9, 14 and 15 after LPS-stimulation was similar to the profiles of the LPS-unstimulated samples. This was further confirmed by the hierarchical clustering performed on the 1000 human genes with the highest mean among samples, and, also in this case, the four samples showed the typical profiles of the LPS-unstimulated ones (data not shown). As we observed differences among samples depending on inter-individual variability to immune response, we tried to obtain more information on the interpersonal reaction to LPS by describing the pattern of expression of a subset of 44 genes that have been previously reported to be a specific signature of induced cytokine response with a heatmap of the sample-to-sample Euclidean distances (36). While in the absence of stimulation no evident differences between samples were observed, after LPS injection, considering these 44 genes two clusters were observed that reflected the traits of the inflammatory response, with high-responders clearly divided from low-responders (Fig. 3b). Of note, since these 44 genes have been identified to be able of deconvoluting complex responses to immune stimulation, we expected the variability between low- and high-responders to be well-defined. Subsequently, we similarly analyzed the Euclidean sample-to-sample distances defined by the expression of all the *hervgdb4* fragments (Fig. 3b). Interestingly, as shown for the 44 immunity genes, also for *hervgdb4* fragments no sample clustering was observed in LPS-unstimulated sample, while in LPS-stimulated samples two clusters were formed, roughly corresponding to low- and high-responders to immune stimulation. However, the three low-responder samples clustering with the high-responder ones do not coincide with the four LPS-stimulated samples that showed a pattern of *hervgdb4* fragments expression similar to the LPS-unstimulated samples in Fig. 3A, suggesting that factors other than the severity of inflammatory response may contribute rather than interfere with the inter-individual variability for *hervgdb4* fragments expression.

**Fig. 2.**
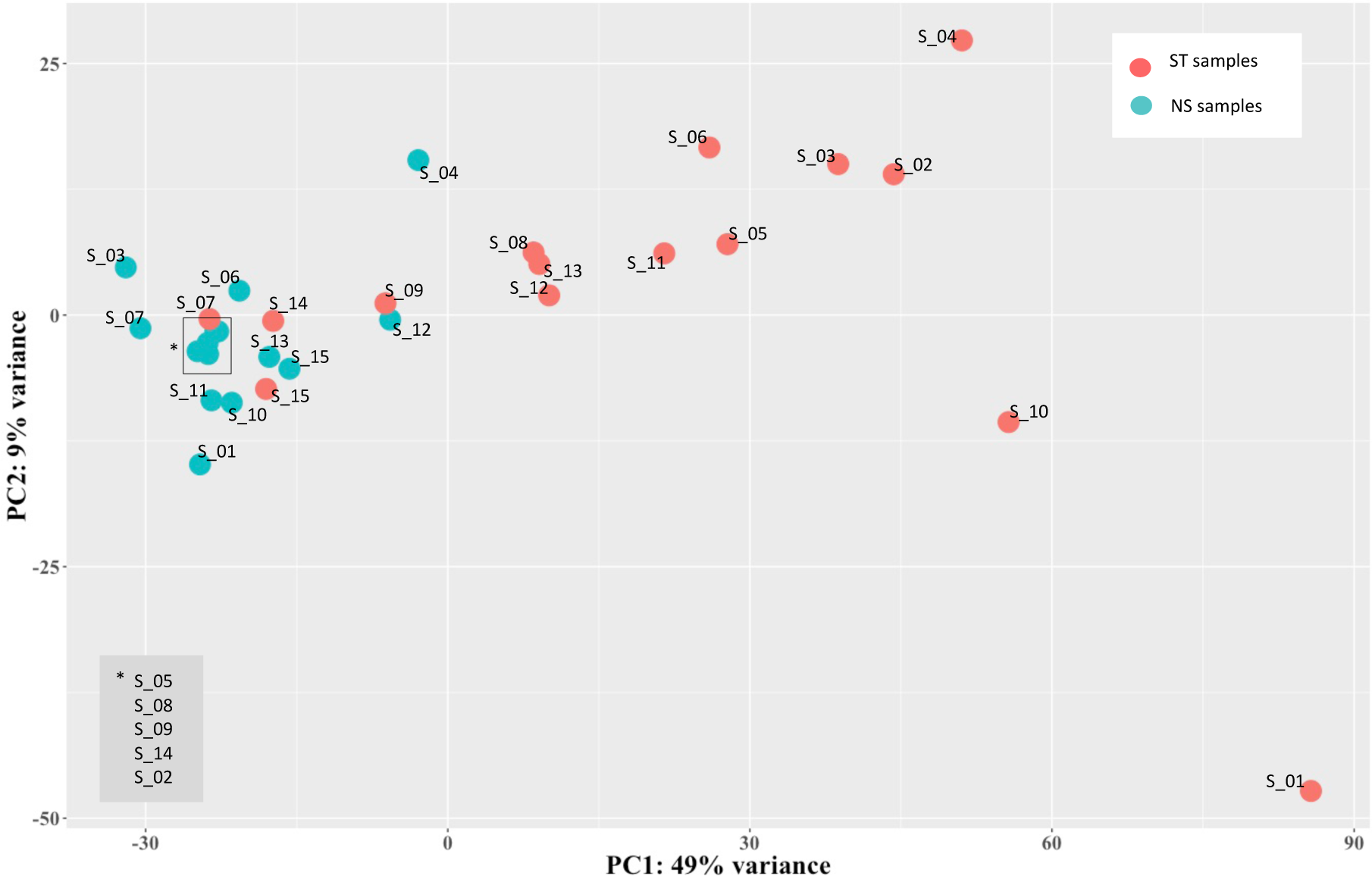
Principal Component Analysis (PCA) of samples. PCA on rlog-normalized *hervgdb4* fragments expression data. It is possible to see the division between not-stimulated and stimulated samples according to the PC1.

**Fig. 3.**
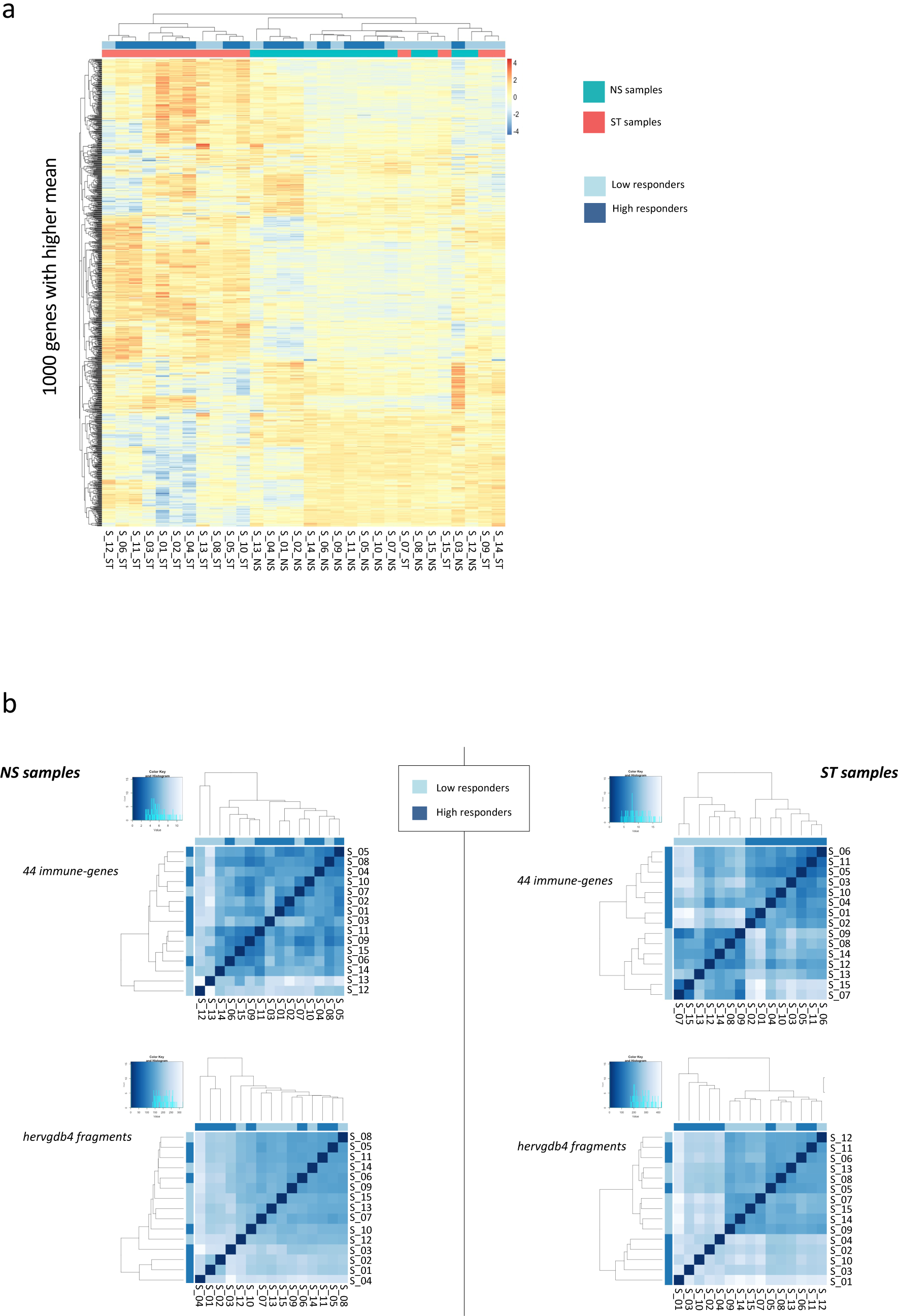
Heatmaps of the overall similarity between samples. (a) Hierarchical clustering of the top 1000 *hervgdb4* fragments with the highest average of rlog-normalized counts, correlation distance measure used in clustering columns. The two clusters highlight specific signatures induced by LPS. (b) Hierarchical clustering of the Euclidean sample-to-sample distance before (NS samples) and after LPS injection (ST samples). We searched for difference related to the pattern of expression of the 44 genes that captured the diversity of complex innate immune responses (36) (44 immune-genes) and of the *hervgdb4* fragments. High- and low-responders showed different response to inflammation.

### Differential LTR-retrotransposon expression in PBMCs

Once assessed that the variability across samples mostly fitted with LPS stimulation, we evaluated the *hervgdb4* fragments for differential expression. After applying a filter (FDR ≤ 0.01 and absolute values of log_2_ Fold Change ≥ 1) we identified a total of 6,452 (11% of the total expressed) Differentially Expressed (DE) *hervgdb4* fragments (Table S1). We represented all the expressed *hervgdb4* fragments in a volcano-plot, where they were indicated as points that spread according to the log_2_ Fold Change on the x-axes and to the adjusted p-value on the y-axes (Fig. 4). It is worth noting that the great majority of *hervgdb4* fragments were positively modulated, showing a general trend of LTR-retrotransposon up-regulation in PBMCs after LPS stimulation. In fact, among the 6,452 DE *hervgdb4* fragments, 5,383(83%) were up-modulated while 1,069 (17%) were down-modulated after stimulation (Table 1). Subsequently, we checked how many *hervgdb4* DE loci were present, observing that 4,607 loci (13% of the total expressed) were modulated. Interestingly, results showed that 3,688 loci (80%) were up-regulated, while 919 loci (20%) were down-regulated. We then focused on the mostly intact ReTe HERV proviruses, observing that 115 of them were DE (17% of the ReTe HERV expressed proviruses). In particular, 86 were up-regulated while 29 were down-regulated. In general, the majority of the DE HERVs was over-expressed: out of the 55 HERV groups, 6 included only up-modulated proviruses, 13 included both up- and down-regulated proviruses and 5 only down-regulated proviruses (Fig. 5). Importantly, we found 23 groups that were constitutively expressed in PBMCs but were not modulated by the stimulation.

**Table 1.** LTR-retrotrasposon modulation. Different proportion of modulated, up- and down-regulated *hervgdb4* fragments (a), *hervgdb4* loci (b) and ReTe proviruses (c) in PBMCs after *in vivo* LPS stimulation

|  | Expressed |  | Modulated |  |
| --- | --- | --- | --- | --- |
|  | Non DE | DE (%*) | Up-modulated | Down-modulated |
| *hervgdb4* fragments | 54,061 | 6,452 (11%) | 5,383 | 1,069 |
| *hervgdb4* loci | 31,079 | 4,607 (13%) | 3,688 | 919 |
| HERV proviruses | 561 | 115 (17%) | 86 | 29 |

\* Proportion of expressed elements that are modulated

**Fig. 4.**
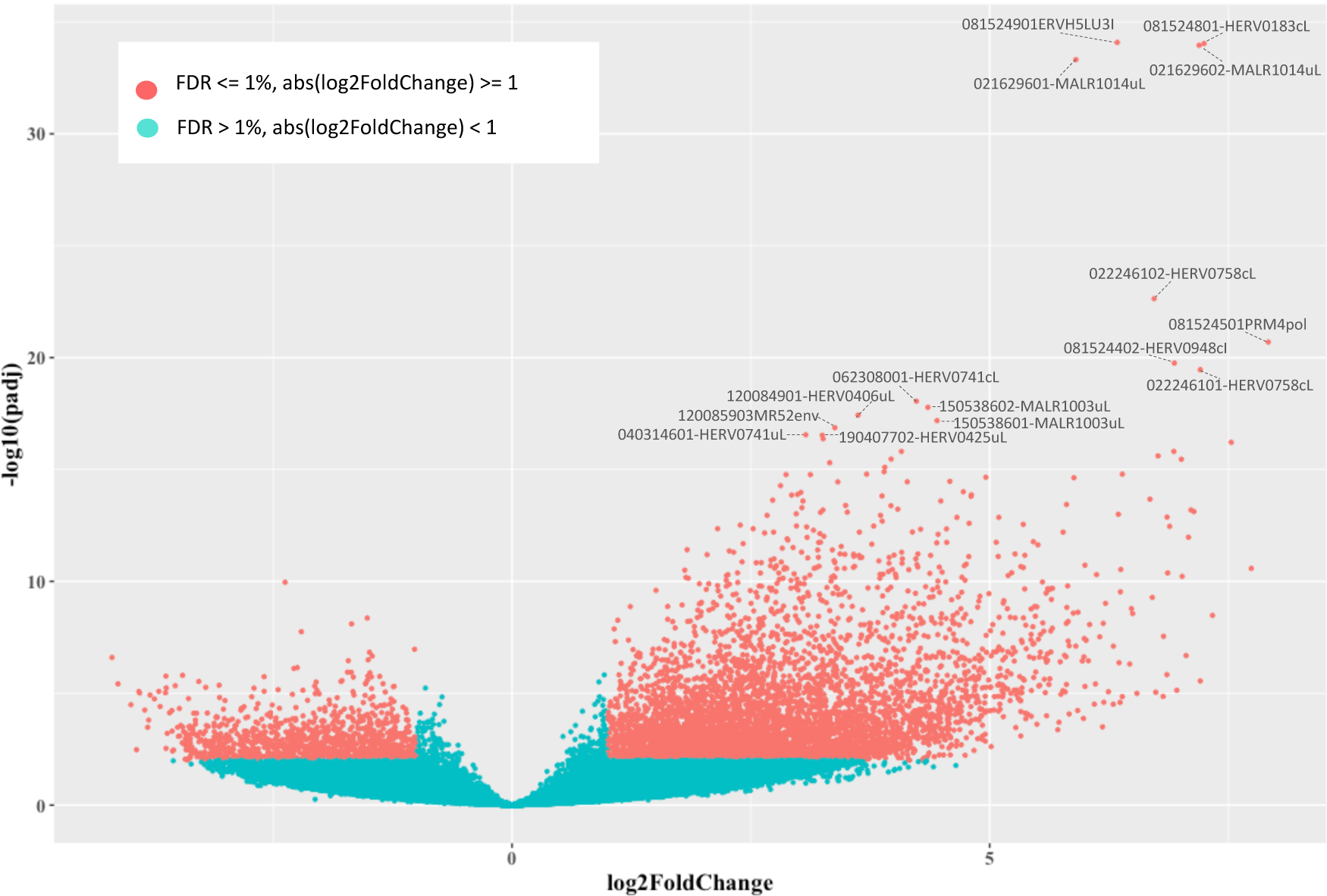
Differential HERV/MaLR expression analysis. Volcano-plot of the differentially expressed *hervgbd4* fragments. Each point represents *hervgbd4* fragments, which spread according to the log_2_ fold change (x-axis), and the log10 adjusted p-values (y-axis). Red points are the significantly modulated *hervgbd4* fragments. For the 15 *hervgbd4* fragments with lowest adjusted p-values, the names are indicated.

**Fig. 5.**
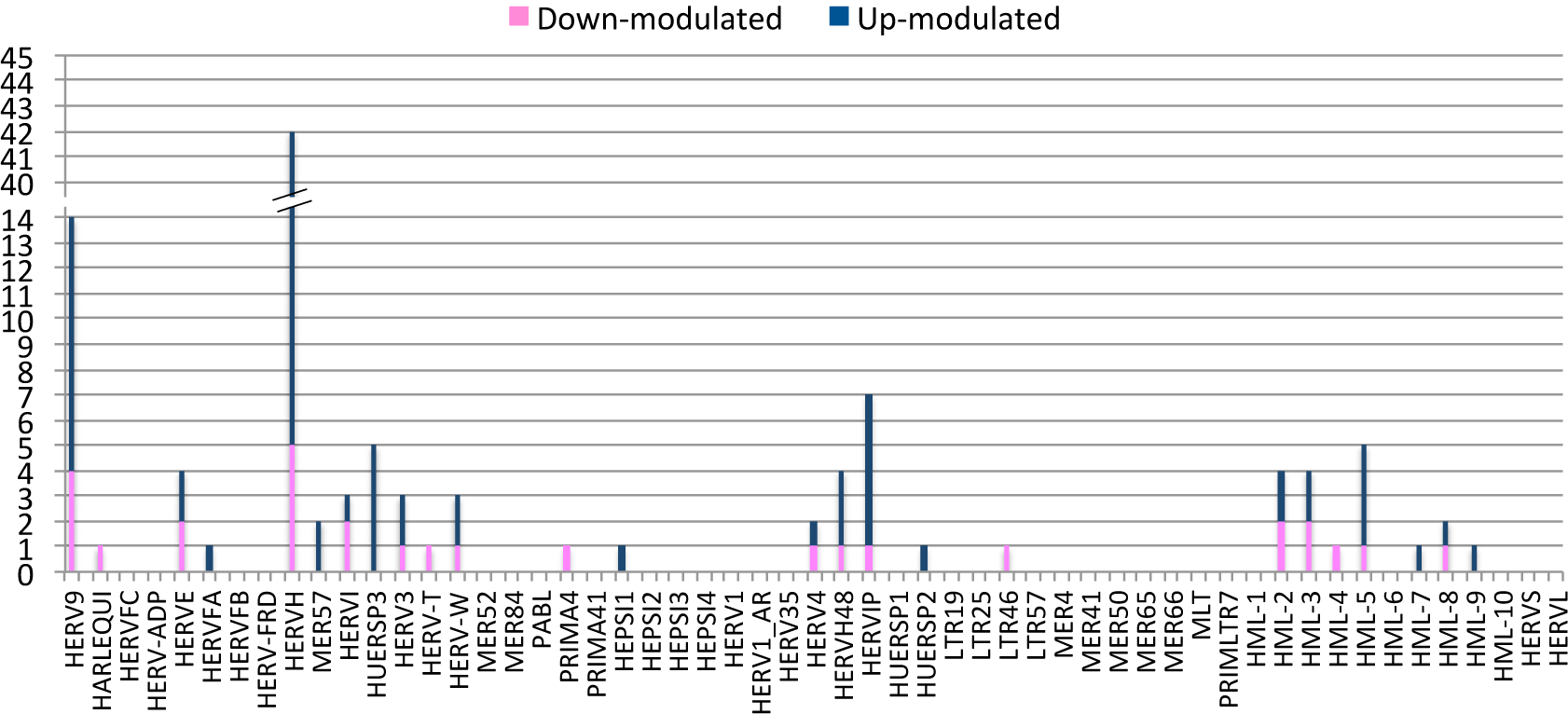
Differentially expressed ReTe HERV proviruses. Bar-plot of the distribution of differentially expressed ReTe HERV proviruses across retroviral classes and groups. Different colors show up- and down-regulation.

### Concordant modulation of LTR-retrotransposons and co-localized immunity-related genes

To gain more insights into the LTR-retrotransposon modulation, we focused on the 15 *hervgdb4* fragments with the highest DE according to their adjusted p-value (Table 2, and Fig. 4). Importantly, all these 15 high DE *hervgdb4* fragments were up-regulated after LPS-stimulation. We used Transcript Per Million (TPM) normalization to have information on the expression levels of the fragments before and after the LPS stimulation (Table 2) and, subsequently, we investigated their context of insertion. Interestingly, we found that, among the 15 most-highly modulated LTR-retrotransposons, 10 were neighbor integrations (within a 10-kb window of distance) with human coding genes. In particular, 6 of them were inside, 3 were downstream and 1 was upstream the colocalized gene. We hence analyzed the modulation of these human genes, observing that all of them were up-regulated as a consequence of LPS stimulation, as summarized in Fig. 6. The fragment with the lowest adjusted p-value (6.99E-36) was 081524901ERVH5LU3I, at coordinates chr8:103002077-103002365. This fragment is the U3 region of a 5’LTR belonging to an HERVH provirus (chr8:103002064-103004587), and its expression levels were increased from an average TPM value of 2.4 to an average TPM value of 124.5 after LPS stimulation. It is worth to note that, even if this LTR sequence is not co-localized with coding genes, it is integrated in a promoter-flanking region that is affected by copy number variation according to ENSEMBL annotations (data not shown), possibly suggesting a potential transcriptional control role. Interestingly, also fragment 081524801-HERV0183cL, part of a solo LTR (chr8:103001306-103001748) within the same region as 081524901ERVH5LU3I, increased its average TPM value from 2.4 to 74.2. Fragments 021629602-MALR1014uL and 021629601-MALR1014uL were part of the same solo LTR at coordinates chr2:113131173-113131620. This solo LTR is integrated within the intron of the Interleukin 1 Receptor Antagonist gene (IL1RA), which codes for a protein that is known to have an anti-inflammatory role (37). Both gene and solo LTR significantly increased their expression levels after LPS administration, showing high average TPM values in stimulated samples (Table 2). Similarly, fragments 022246101-HERV0758cL and 022246102-HERV0758cL, part of the same solo LTR at coordinates chr2:113131173-113131620, showed a pattern of up-regulation comparable with their neighbor gene, namely TNF alpha induced protein 6 (TNFAIP6). Thus, also in this case, the solo LTR is co-localized with a gene that is involved in immunity, having a known regulatory function (38). Instead, fragments 081524501PRM4pol and 081524402-HERV0948cI, which are portions of the internal regions in proviral loci chr8:102984015-102986529 and chr8:102980120-102991770, respectively, were found to be intergenic integration. The basal expression levels of both fragments were 0.1 TMP, increasing to 3.6 and 5.5 TPM after stimulation, respectively. Next, we found that fragment 062308001-HERV0741cL, a solo LTR in chr6:159677828-159678891, is integrated within the intron of Superoxide dismutase 2 (SOD2) gene. Of note, fragments 150538602-MALR1003uL and 150538601-MALR1003uL, both part of a solo LTR in chr15:63906995-63907370, showed an average TPM increased from 3.7 and 1.0 to 49.7 and 14.9 after stimulation, respectively, are integrated within the 3’ untranslated region (UTR) of the Death-associated Protein Kinase 2 (DAPK2) gene. Fragment 120084901-HERV0406uL, a solo LTR in locus chr12:8537686-8538696, is integrated inside the C-type lectin domain family 4 member E (CLEC4E) gene, and increased its average TPM values from 1.0 to 87.2. The fragments 120085903MR52env (chr12:8566648-8568185), representing an intergenic integration, and 040314601-HERV0741uL (chr4:15677257-15678021), being inserted upstream the Family With Sequence Similarity 200 Member B gene (FAM200B), showed more than 5-folds increase in average TPM after LPS stimulation. Finally, the fragment at coordinates chr19:35887015-35887532, which increased its average TPM values from 5.7 to 28.9, is integrated downstream the NFKB inhibitor delta gene.

**Table 2.**
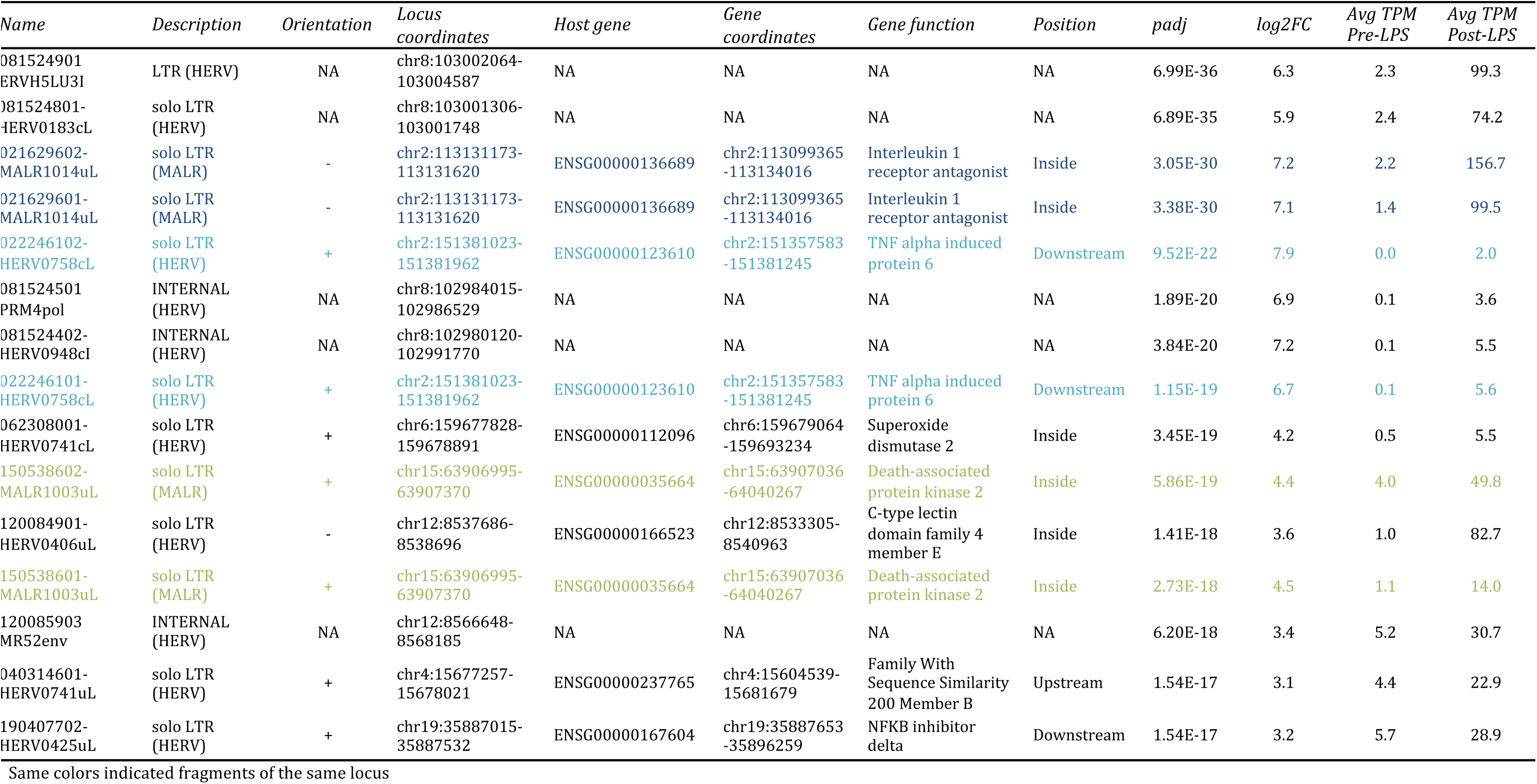
Top 15 most DE *hervgdb4* fragments. Description of the most 15 most DE *hervgdb4* fragments, including their expression levels indicated as TPM values, and their context of insertion.

**Fig. 6.**
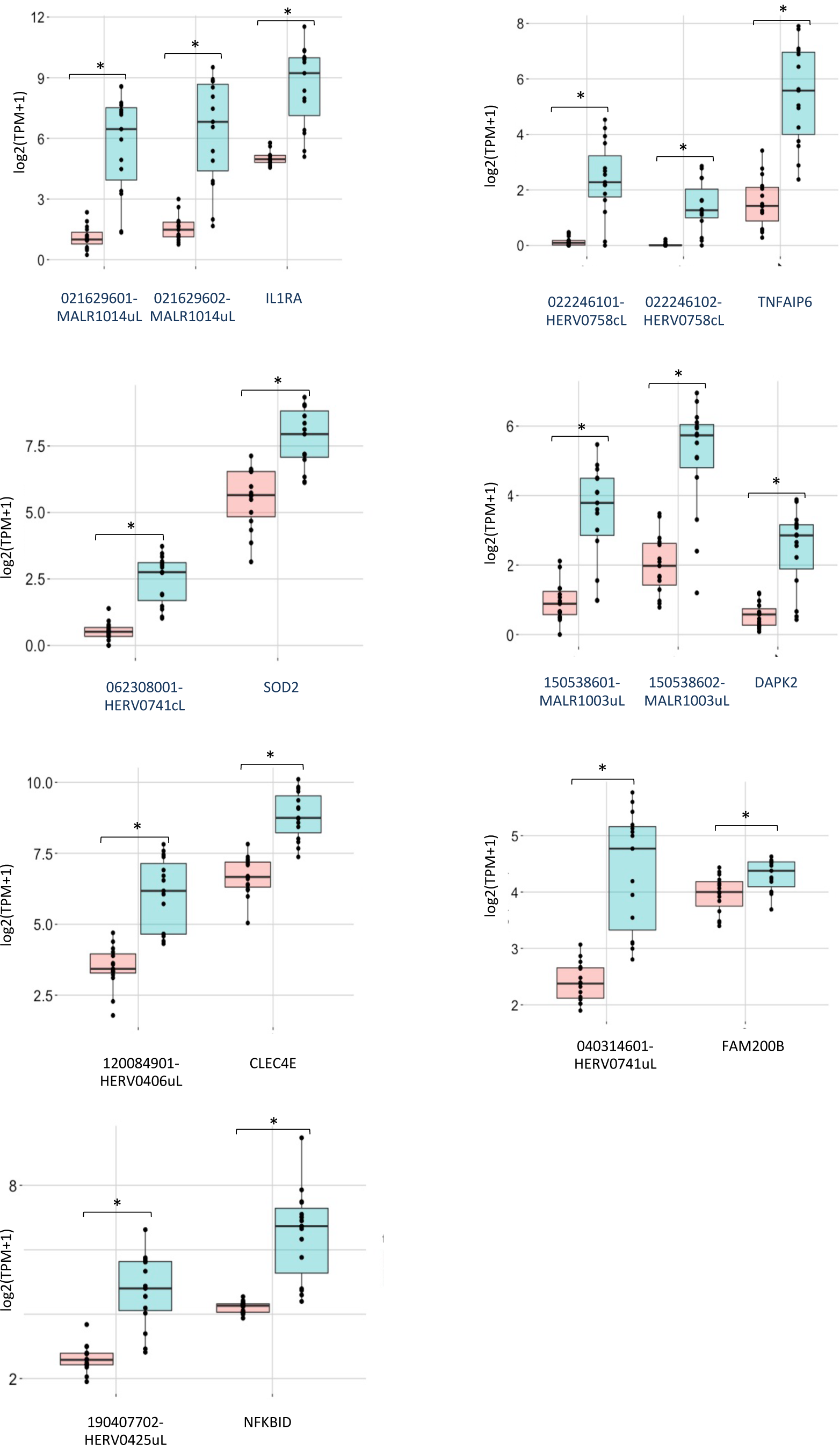
Co-regulation of *hervgbd4* fragments and human genes. Boxplot of the TMP expression values of the *hervgbd4* fragments co-localized with human genes and the neighbor genes. The red boxes indicated values from non-stimulated samples; blue boxes indicated values from stimulated samples. Significant modulations according to the DEseq2 analysis (padj < 0.01) are marked with stars.

## Discussion

We used an RNA-seq approach and the hervgdb4 database (33) to assess an overview of the LTR-retrotransposons transcriptome in PBMCs. This analysis has allowed, for the first time, to investigate HERVs and MaLRs proviruses and solo LTRs expression in human patients by mimicking gram-negative bacterial infection, reveling a total of 35686 LTR-retrotransposon elements expressed in PBMCs.

Previously, the same database has been used by Mommert *et al*. (29) to identify expressed and modulated loci in an *ex vivo* system of LPS stimulation and endotoxin tolerance, through microarray analyses (29). Interestingly, the percentage of expressed elements in our *in vivo* model (6.8% of *hervgdb4* fragments and 8.4% of *hervgdb4* loci expressed) was slightly higher than the one measured in *in vitro* experiments (5.6% of *hervgdb4* fragments). Among all groups, class II groups are those most active, while groups belonging to class III were, in general, less active. In particular, HML-2 is the most expressed in PBMCs. This group is also one of the most investigated, due to recent HML-2 integrations and the hypothesized implication of some of active loci in several diseases (39–42). In comparison to Mommert *et al*. (29), who reported instead an abundant activation of several class I and of all class III groups, the LTR-retrotrasposon expression among classes showed some differences. These differences can be explained by i) the technologies and methods that have been used (*in vivo* vs *ex vivo*, microarray vs RNA-seq), and ii) the great differences in the basal transcriptional activity of each individual, which have been already observed in PBMCs (43). Moreover, the use of the Vargiu *et al*. database (4) has allowed us to analyze the expression of the most intact HERV proviruses for which we can hypothesize a higher probability of protein production. The pattern of counts distribution among classes of the most intact HERV proviruses is similar to the one obtained when considering the *hervgdb4* loci, and it is mostly in agreement with the information on HERV-H, HML-2, HERV-E and HERV-W transcriptional activity in PBMCs already reported in previous studies (37). Importantly, this study has given information on the overall expression of the groups, but not on the single loci. Moreover, the expression of HERV-E has been reported to be characteristic of only a small percentage of the subjects analyzed, while a large portion of both *hervgdb4* loci (the 24% of the group) and most intact HERVs (the 28% of the group) is expressed in PBMCs, in the present study.

The analysis of the variability among the 30 samples analyzed showed differences between non-stimulated samples and LPS-stimulated samples. Indeed, in both PCA analysis and hierarchical clustering the not-stimulated and stimulated samples spread in two distinct clusters. However, a great interpersonal variability on the response of patients to LPS is also evident. This heterogeneity in the HERVs and MaLRs expression, as a consequence of LPS stimulus, is in line with the already observed strong inter-individual variability of gene expression in response to microbial agents (36, 44). Importantly, the definition of such variability should be a necessary foundation for better understanding a possible implication of HERVs and MaLRs expression in immunity and disease.

The analysis of *in vivo* HERVs and MaLRs modulation reveals a total of 6,452 hervgdb4 fragments modulated in PBMCs, with a clear trend of up-regulation induced by LPS response (81%). In comparison to Mommert *et al*. (29), in which 111 DE HERVs and MaLRs (44 down-regulated and 67 up-regulated) have been identified, a larger number of DE elements was observed, which can be explain by the different sensitivity of the methods used. Specifically, only 18 of the DE *hervgdb4* fragments identified by Mommert *et al*. are included in the present list of DE *hervgdb4* fragments, suggesting that the two models are not equal but complementary. Moreover, another study has identified a list of LTRs-retrotransposon INF-induced (45) and a proportion of them (about the 16%) was observed in this study to be also LPS-induced (data not shown). This highlights the importance of investigating the LTRs-retrotransposon modulation in several conditions, but it also suggests that distinctive patterns of LTRs-retrotransposon induction, if used as biomarker, may became indicative to separate physiological from pathological conditions. The LPS stimulus also provoked a strong activation of the most intact ReTe proviruses. Importantly, not all the groups respond in the same way to the LPS stimulation.

Among the 15 most modulated hervgdb4 fragments, 10 were co-localized with human genes. We used the average expression level of the most DE human gene, IL1A (ENSG00000115008), which showed an average TMP of 19.4 after LPS stimulation (data not shown), as a comparison to understand a possible biological meaning of the expression of the fragments we found modulated. In general, all the fragments are not only modulated but also generally medium/highly expressed. Among 12 *hervgdb4* loci, 10 are LTRs, including 9 solo LTRs. This kind of information might be integrated by other studies on transcription factor binding sites (TFBS) provided by HERVs and MaLRs (14), including those for NF-kB, IRFs and STATs, suggesting a possible HERVs and MaLRs activation in immunity (46, 47). Moreover, 10 *hervgdb4* fragments are neighbor integrations of human genes that are also activated after inflammatory response. Of particular interest is the identification of 3 solo LTRs localized outside the modulated genes, due to the possible presence of promoters or polyadenylation signals that may play a role in the regulation of the nearby gene. Specifically, the solo LTRs in chr2:151381023-151381962 and chr19:35887015-35887532 are integrated downstream the TNF alpha induced protein 6 gene (TNFAIP6) and NF-kappa-B inhibitor delta (NFKBID), respectively. Interestingly, while the most DE genes identified in this model are mainly those coding for proteins that are positive regulators of immunity, such as IL1A and IL1B (data not shown), these data showed a strong up-regulation of LTRs-retrotransposon co-localized with genes coding for cellular inhibitors of these proteins. Hence, if on the one hand it has been suggested that a subset of HERVs hold TFBSs (47) that may increase their activation in immunity, on the other hand the genes products of TNFAIP6 and NFKBID are potentially able to inhibit this phenomena(48, 49). For this reason, this data underlines the complexity of the relationship between the HERVs/MalRs modulation and the immune response, especially if hypothesizing their active role on the regulation of the co-localized immune-related genes. Certainly, this data may give the bases to clarify the LTRs-retrotransposon involvement in the regulation of immune functions, but further studies are needed to clarify these mechanisms and may help to understand how HERVs and MaLRs contribute to the complexity of immune response.

## Conclusions

We used RNA-seq transcriptome data from 15 healthy participants to a clinical trial injected with LPS, with the aim to identify expressed and modulated LTR-retrotransposon elements. This RNA-seq based approach revealed the basal expression of 18,633 HERV *hervgdb4* loci, including those of 723 most intact ReTe proviruses, and 17,053 MaLR *hervgdb4* loci, in PBMCs. To assess the specific signatures induced by LPS we investigated the sample-to sample differences, which showed a specific pattern of *hervgdb4* fragments expression as a consequence of inflammation. We hence evaluated the *hervgdb4* fragments modulation, identifying a total of 6,452 differentially expressed elements. Moreover, we observed a general trend of activation of the LTR-retrotransposons after LPS stimulation. Interestingly, the HERVs/MaLRs regulation was similar to the cellular gens regulation, suggesting possible interactions between LTR-retrotransposons and the immune response. However, further analyses are required to evaluate whether such modulation is involved in the regulation of immune response and to better assess the LTR-retrotransposons expression significance in immunity.

## Methods

### RNA-seq data and quality control

We used a RNA-seq dataset public available (GEO:GSE87290), including the transcriptome of PBMCs from healthy humans (n=15) before and after 1ng/kg LPS exposure. We checked the quality of the RNA sequences by using FastQC Galaxy Version 0.72 (http://www.bioinformatics.babraham.ac.uk/projects/fastqc). Low quality reads were trimmed with Trim Galore! V.0.4.3.1 (https://github.com/FelixKrueger/TrimGalore).

All the mentioned analyses were done on Galaxy release_17.09 locally installed (http://galaxyproject.org/).

### Reads mapping and counting

HISAT2 Galaxy Version 2.1.0 was used with default parameters to map reads to genome assembly hg38. We assessed the quality of the alignments by using the stats function of bamtools Galaxy Version 2.0.1 (50). We counted the reads mapping to each *hervgdb4* fragment and human gene by using the “union” mode in htseq-count Galaxy Version 0.6.1galaxy3 (51), and *hervgdb4* database (33) and gencode.v27 (52) for respectively LTR-retrotransposon and gene annotations. All the mentioned analyses were done on Galaxy release_17.09 locally installed (http://galaxyproject.org/). We calculated the expression values of the expressed *hervgdb4* fragments and genes as Transcripts Per Million (TPM).

### Raw data filtering and transcriptome analysis

We selected all the genes and *hervgdb4* fragments with at least 1 count in at least 10 samples, conventionally considering them as expressed in at least one experimental condition and we used the obtained data for the analysis of HERVs/MaLRs transcriptome in PBMCs. We used the GenomicRanges v.1.30.3 (53) R (version 3.4.4) package to obtain the coordinates of the most intact HERV proviruses (4) from those of the expressed *hervgdb4* fragments.

### Hierarchical clustering

The DESeq2.v.1.18.1 R (version 3.4.4) package (54) has been used to perform rlog normalization on human genes and HERVs/MaLRs raw counts. From the output of the normalization we extracted the HERVs/MaLRs rlog counts and we assessed the interpersonal variability through PCA and Heatmap. The PCA was built with the function plotPCA in DESeq2.v.1.18.1 and visualized by using ggplot 3.0.0 in R (version 3.4.4). The Heatmap of the 1000 *hervgdb4* fragments with the higher average rlog counts across samples was built through the pheatmap 1.0.10 R (version 3.4.4) package, considering the correlation distance across samples in column. The dist function in DESeq2.v.1.18.1 was applied to the transpose of the rlog transformed count matrix to get Euclidean sample-to-sample distances; the heatmaps were built through the pheatmap 1.0.10 R (version 3.4.4) package.

### Differential gene and *hervgdb4* fragment expression analysis

We performed a differential expression analysis of the genes and *hervgdb4* fragments by using the DESeq2.v.1.18.1 R (version 3.4.4) package (54) and the filtered raw counts as input. During the analysis, false discovery rate/adjusted p-value were used for multiple test comparison according to the Benjamini-Hochberg procedure (55). We used a threshold (FDR <= 0.01 and absolute values of log_2_ Fold Change >= 1) to identify the modulated elements. The differentially expressed *hervgdb4* fragments were visualized in a volcano plot built by using ggplot 3.0.0 in R (version 3.4.4). We sorted the *hervgdb4* fragments by adjusted p-values to recognize the mostly differentially expressed. We compared the TPM expression values with the filtered adjusted p-values (<= 0.01) to see the relative distributions, as describe in Fig. S1. We manually checked for the presence of neighbor genes on ENSEMBL (56), within a 10-kb window of distance from the nearest-neighbor gene.

---

**Fig. S1.**
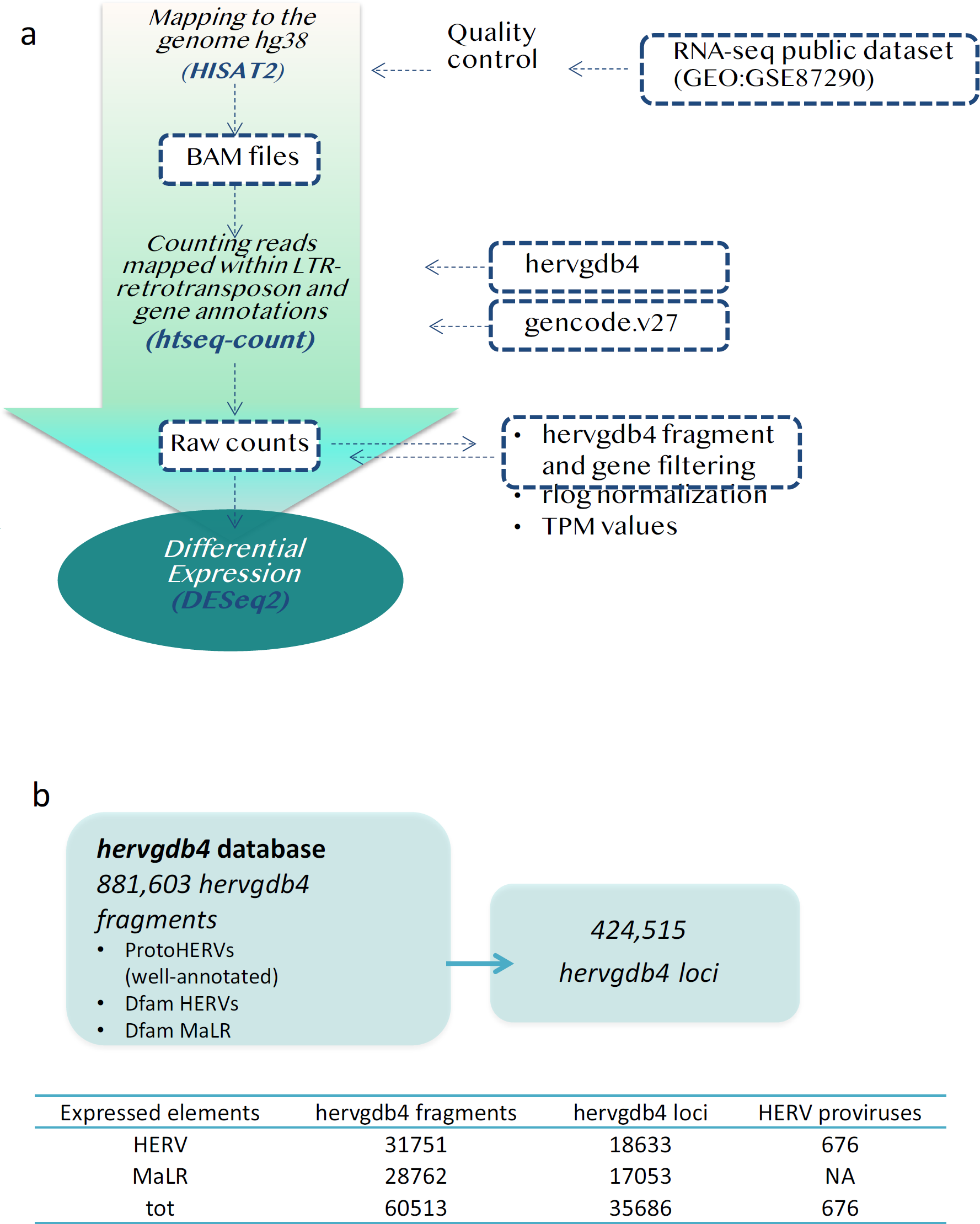
Experimental design of Differential Expression analysis. RNA-seq workflow for the identification of modulated HERVs and MalRs (a). The input files used are in blue boxes. The composition of hervgdb4 database is schematized in (b). The amount of expressed hervgdb4 fragments and loci have been obtained by filtering the raw counts, and are summarized in the table. The coordinates of the expressed ReTE most intact proviruses have been obtained by using the findOverlaps function from package “Iranges” and the coordinates of expressed hervgdb4 fragments.

